# Huntingtin bundles and changes the local proteome of actin filaments in neurons

**DOI:** 10.1101/2023.08.29.555277

**Authors:** Rémi Carpentier, Mariacristina Capizzi, Hyeongju Kim, Julia Novion-Ducassou, Eric Denarier, Béatrice Blot, Yohann Couté, Isabelle Arnal, Ji-Joon Song, Sandrine Humbert

**Affiliations:** Univ. Grenoble Alpes, Inserm, U1216, CEA, Grenoble Institute Neurosciences, 38000 Grenoble, France; Institut du Cerveau-Paris Brain Institute, Sorbonne Université, Inserm, CNRS, Hôpital Pitié-Salpêtrière, Paris, France; Department of Biological Sciences, Korea Advanced Institute of Science and Technology (KAIST), Daejeon 31414, Korea; Univ. Grenoble Alpes, INSERM, CEA, UMR BioSanté U1292, CNRS, CEA, FR2048, 38000 Grenoble, France

## Abstract

Huntingtin (HTT) is a large protein whose best-known function being the facilitation of intracellular dynamics along the microtubule network by scaffolding molecular motors complexes. Our recent finding that the defective axonal growth in HD was due to altered growth cone architecture led us to ask whether HTT also influences the cytoskeleton itself. In developing neurons, we found that a large proportion of HTT associates with F-actin in growth cones. Using cell free system and purified recombinant proteins, we observed that HTT binds directly filamentous actin (F-actin) and organizes filaments into bundles. Transmission electron microscopy shows that HTT dimers crosslink adjacent filaments 20 nm apart. We also provide evidence that HTT binding on F-actin modulates the association of other proteins to this cytoskeleton. Notably, HTT limits the association of the growth cone protein Drebrin1 with F-actin. HTT depletion leads to abnormal cytoskeletal organization, localization of Drebrin1 in growth cones, and axonal growth. HTT therefore serves a scaffolding function for the cytoskeleton itself, what might be relevant for HD pathophysiology.

## Introduction

The Huntingtin (HTT) protein was discovered because its N-terminal polyglutamine repeat is abnormally expanded in Huntington’s disease (HD) (Group 1993). HTT is ubiquitously expressed, but mutant HTT (mHTT) tends to wreak the most havoc in the brain, causing neuronal dysfunction and loss in the cortex and in the striatum that manifest in psychiatric, cognitive, and motor deficits by mid-adulthood. HTT is a flexible protein that can adopt a hundred distinct conformations, with multiple HEAT domains that enable it to scaffold numerous proteins and serve in cellular functions from gene expression to signal transduction, cell metabolism, and cellular dynamics (Saudou and Humbert 2016). At a molecular level, HTT is best known, however, for its roles in intracellular transport and vesicular recycling. HTT controls the transport directionality of vesicles and organelles, depending on its phosphorylation state, which determines whether it scaffolds its molecular cargo and various accessory proteins with dynein or kinesin motors (McGuire et al. 2006; Caviston et al. 2007; Twelvetrees et al. 2010). HTT participates to numerous cellular processes during cortical development by controlling the transport of proteins to precise subcellular localization. HTT controls the shuttling of PCM-1 between the cytoplasm and the base of the cilium during ciliogenesis (Keryer et al. 2011), the targeting of Dynein and NUMA1 to the cell cortex of cortical progenitors to regulate the orientation of the mitotic spindle (Godin et al. 2010), and the recycling of N-cadherin molecules during the radial migration of new-born neurons (Barnat et al. 2017).

While it is clear that HTT relies on microtubules for many of its activities, a recent discovery raised the possibility that HTT might be involved in organizing the microtubules themselves. We had been pursuing the developmental origins of cortical defects in HD and traced the growth of callosal projection neurons in developing mice (Capizzi et al. 2022). The axons in HD mice were shorter than in wild-type animals—many axons did not even cross the callosum, and arborization on the contralateral side was sparser, resulting in weaker connectivity between the two hemispheres. This axonal outgrowth defect was due to the disruption of the microtubule organization of growth cones, the swellings at the axon tips that respond to attractive and repulsive chemical guidance signals and cause the axon to grow through the interplay of actin filaments and microtubules (Dent et al. 2011; Miller and Suter 2018; Pinto-Costa and Sousa 2021; Alfadil and Bradke 2022; Atkins et al. 2022). This finding shows that the defect in microtubule-based vesicular transport observed in HD may not only arise from alteration of the HTT scaffolding function of the molecular motors, but also from disorganization of the microtubules roads themselves. This raised many new questions, such as whether this occurs only in the context of HD, and whether HTT might also influence the structure of the actin cytoskeleton.

There have been previous indications that HTT was at least indirectly involved with actin. In fact, a few studies have reported that when HTT is absent or mutated, actin dynamics are altered (Lynch et al. 2007; Munsie et al. 2011; Tourette et al. 2014). Bioinformatic analyses of the numerous HTT primary and secondary interactors suggest that HTT is involved with Rho GTPase signaling (Tourette et al. 2014), which transduces extracellular signals into coordinated changes in the actin cytoskeleton; in fact, HTT modulates the activity of the small GTPase Rac1 (Tousley et al. 2019b) and interacts with both its Rho guanine nucleotide exchange factor Kalirin (McClory et al. 2018) and its adaptor protein IRSp53 (Tourette et al. 2014). Proteomic, immunoprecipitation and proximity ligation studies have shown that HTT interacts with several actin-binding proteins, such as the G-actin binding protein Profilin, the actin bundling protein α-actinin, the molecular motor Myosin Va or the Huntingtin Interacting Proteins HIP1/HIP1R (Culver et al. 2012; Shirasaki et al. 2012; Tourette et al. 2014; Tousley et al. 2019a). HTT has been also observed to colocalize with several actin structures such as lamellipodia, stress fibers, and nuclear actin rods (Munsie et al. 2011; Miller et al. 2012; Tousley et al. 2019a)(Miller *et al*., 2012; Munsie *et al*., 2011; Tousley *et al*., 2019a) and a previous studies reported that a short fragment of HTT co-sediments with F-actin (Angeli et al. 2010).

To investigate HTT’s possible direct functions on the actin cytoskeleton, we employed cell-free system approaches using full-length human recombinant HTT, proteomic analysis on *in vitro* reconstituted systems and cultured cortical neurons in which the *huntingtin* gene can be specifically depleted.

## Results

### HTT associates with F-actin in growth cones’ filopodia

Together with the fact that we previously showed an axonal outgrowth defect in HD due to the disruption of the microtubule array of growth cones (Capizzi et al. 2022), we explore the HTT possible functions on the growth cone’s cytoskeleton machinery. We first verified that HTT was expressed in growth cones during the developmental axonal outgrowth. Cortical neurons extend short undifferentiated neurites until two days *in vitro* (DIV2), at which point one neurite becomes elongated; this long process becomes the axon and undergoes fast outgrowth during the following DIV, while dendrites will start to elongate and branch (Polleux and Snider 2010; Alfadil and Bradke 2022). Western blot showed that HTT expression doubled between DIV1 and DIV4 and then stabilized (Supplementary Fig. 1). We assessed the subcellular localization of HTT by immunolabelling cortical neurons at DIV4 for HTT and the axon marker Tau-1 (Fig. 1A). HTT was visible in the nucleus, the cell body, along the axonal shaft and in the axonal growth cones. To confirm these observations, we used a sucrose gradient fractionation protocol to isolate growth cones from extracts of developing cortices (Leshchyns’ka and Sytnyk 2013). As expected, the growth cone marker JNK-interacting protein 1 (JIP1) was present in the growth cone fraction, while the non-growth cone marker cis-Golgi matrix protein 130 (GM130) was present in the non-growth cone fraction that contains the soma and nuclei. HTT was present in both fractions (Fig. 1B).

**Figure 1.**
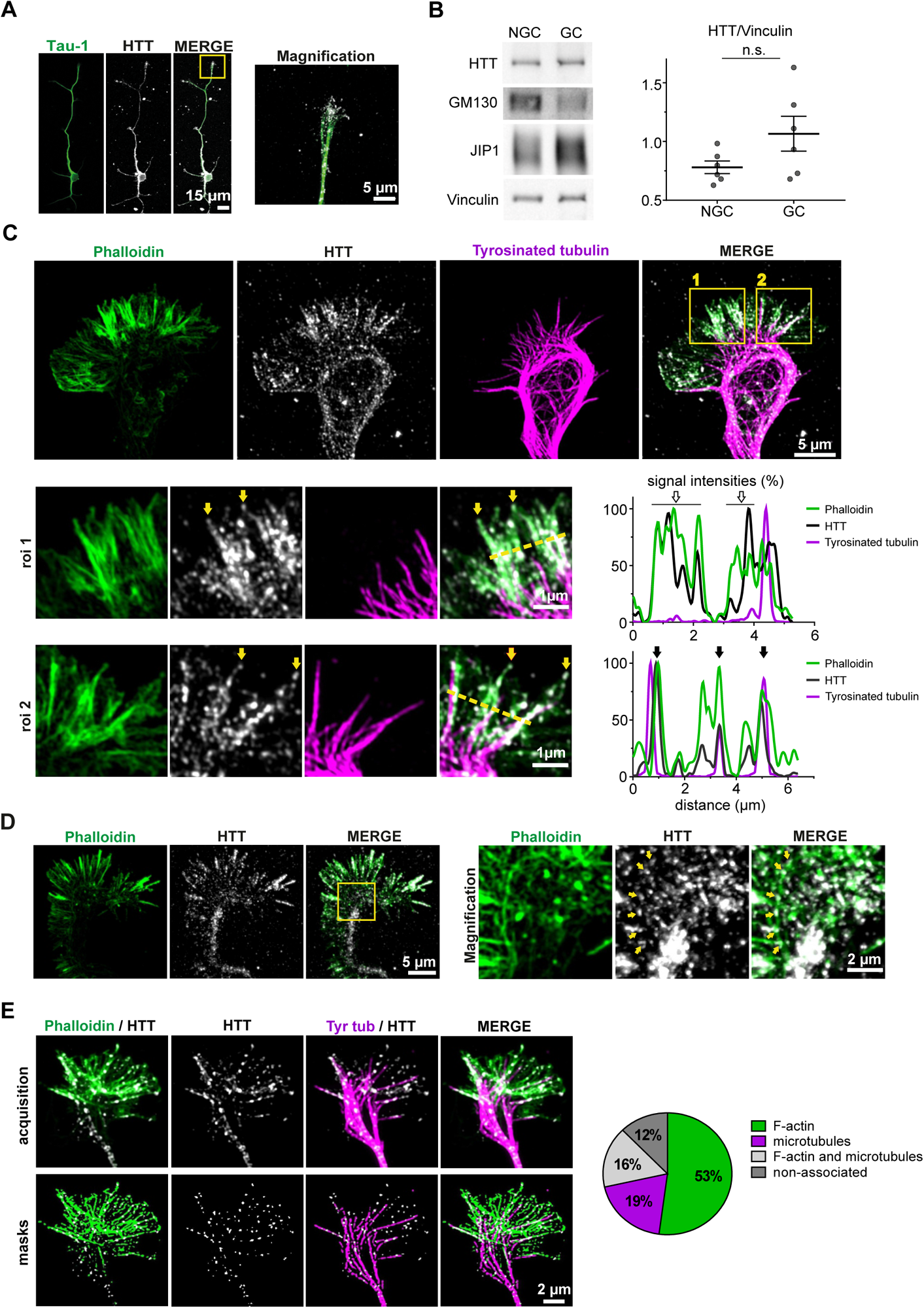
HTT localizes with actin and microtubule cytoskeletons in axonal growth cones. (**A**) Confocal images of a neuron immunostained for the axonal marker Tau-1 (green) and endogenous HTT (grey). Magnification of the region indicated by the red box shows the growth cone. (**B**) Left: Representative western blot of growth cone (GC) and non-growth cone (NGC) fractions. Right: Quantification of HTT protein levels normalized to Vinculin in growth cone and non-growth cone fractions; not significant (n.s.). Paired t-test (NGC: 0.78 ±0.13; GC: 1.07 ±0.34; n=6 fractionations). Data are presented as mean ±SD. (**C**) Airyscan confocal image of a DIV4 growth cone immunostained for F-actin (phalloidin, green), endogenous HTT (grey), and tyrosinated tubulin (magenta). Areas marked by yellow boxes are magnified in rows 1 and 2. Magnification 1 shows that HTT localizes with F-actin in filopodia in the absence of exploratory microtubules. Magnification 2 shows that HTT localizes at the interface of F-actin and exploratory microtubules. Yellow arrowheads mark HTT localizing at the tip of filopodia and yellow dotted lines indicate the axis along which the representative line-scan analyses were performed. Bottom right: Representative line-scan analyses (relative fluorescence intensity) of indicated immunostainings. Empty arrows indicate phalloidin/HTT co-localization and black arrows indicate regions of phalloidin/HTT/Tyrosinated tubulin triple-co-localization. (**D**) Left: representative airyscan confocal acquisition of wild-type growth cone labeled for F-actin (phalloidin, green) and HTT (4C8, grey) showing localization of HTT on actin arcs, the yellow box shows the magnified area. Right: Magnified area, yellow arrows show HTT localizing on actin arcs. (**E**) Representative confocal images (top) and associated masks (bottom) obtained with macro analysis of a growth cone of DIV4 neuron immunostained for F-actin (phalloidin, green), HTT (grey) and Tyrosinated tubulin (Tyr tub, magenta). Pie chart to the right displays the proportion of HTT 3D objects localizing with F-actin, microtubules, F-actin and microtubules, and the ones non-associated to any cytoskeletons; n=21 growth cones.

Growth cones are composed of three regions. The distal end is the peripheral domain, where the axon grows by extending actin-rich filopodia (spike-like cylinders of parallel F-actin bundles), lamellipodia (flat webs of branched F-actin), and some exploratory microtubules. Some anti-parallel F-actin bundles form contractile arcs into the transitional domain. The central domain, which stabilizes the axon shaft, consists of bundled microtubules whose plus-ends face the transitional domain. To determine where HTT localizes relative to the cytoskeletal components of the growth cone, we labeled F-actin with phalloidin and used antibodies directed against HTT and tyrosinated-tubulin to visualize dynamic microtubules (Fig. 1C). HTT colocalized with both microtubules and F-actin throughout the growth cone: we observed endogenous HTT to be particularly enriched with F-actin bundles in filopodia and along the interface between F-actin and exploratory microtubules invading filopodia (Fig. 1C), while HTT staining punctually localizes at actin arcs in the transition domain without particular enrichment (Fig. 1D, yellow arrows). To understand how HTT distributes along the actin and microtubule cytoskeletons in growth cones, we used high-resolution Airyscan imaging. We created masks based on the signals from HTT, F-actin and microtubule staining and calculated the proportion of HTT-positive objects localizing with F-actin and microtubules (Fig. 1E). Approximately 50% of HTT co-localized with F-actin, 20% with microtubules and 16% overlapped with both networks. Only about 12% of HTT was not associated with the cytoskeletal network.

To confirm that endogenous HTT associates with the cytoskeleton, we used a detergent extraction approach to remove cytosolic proteins while allowing cytoskeletal proteins to remain attached (Fassier et al. 2018). Whereas permeabilization decreased the mean fluorescence intensity of cytosolic and exogenously expressed fluorescent protein Venus within growth cones (Fig. 2A and 2B), the endogenous HTT remained colocalized with microtubules and F-actin. Next, we incubated pre-polymerized microtubules or F-actin with lysates of developing cortices and performed high-speed ultracentrifugation to see what proteins co-sedimented with polymerized cytoskeletons (Fig. 2C and 2D). As a control, we used a sample without any polymerized cytoskeleton to evaluate the amount of HTT present in the pellet as a result of aggregation. The endogenous HTT co-sedimented with microtubules, as previously reported (Gutekunst et al. 1995; Tukamoto et al. 1997) but it also co-sedimented with F-actin.

**Figure 2.**
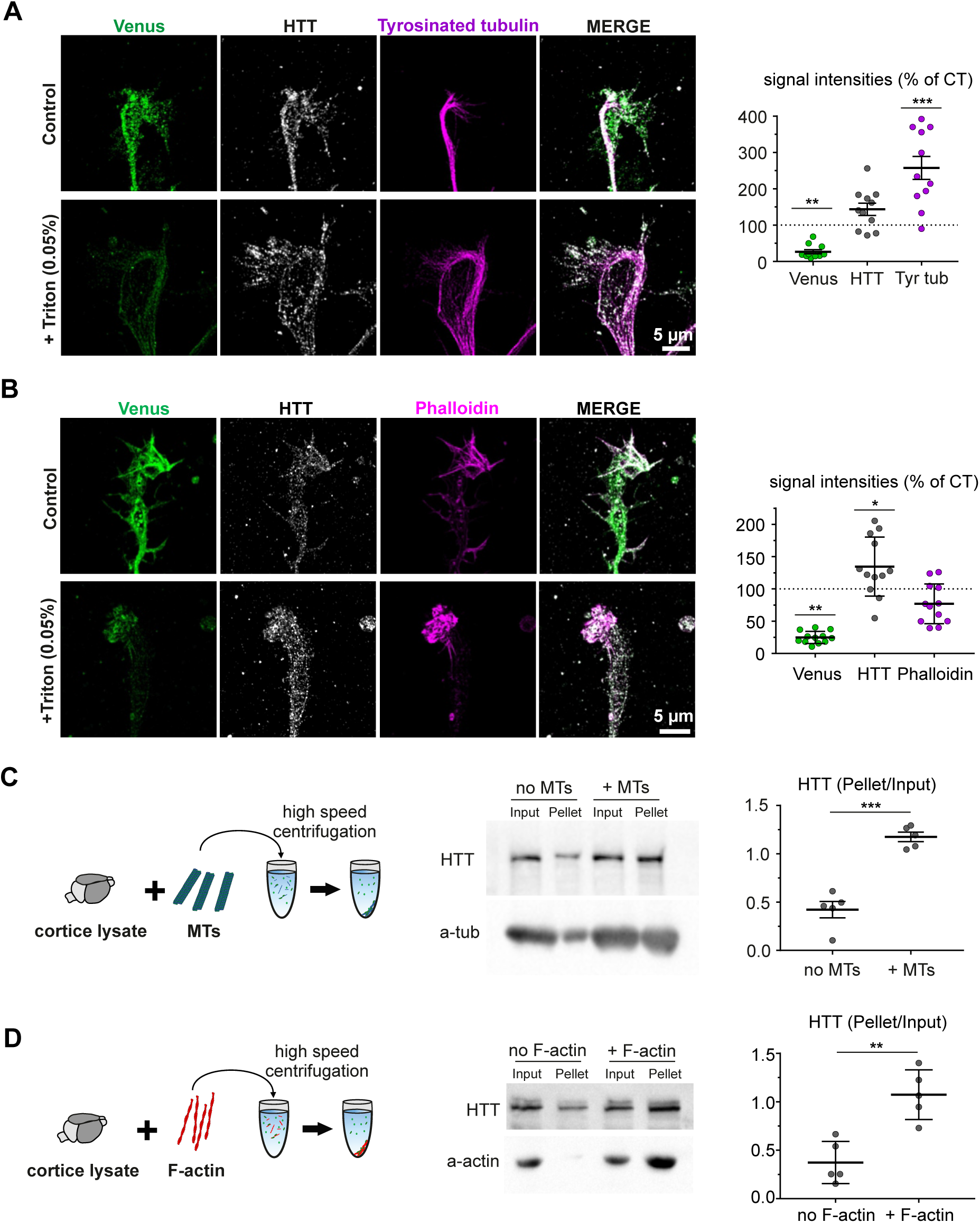
HTT associates with microtubules and F-actin in growth cones. (**A**) *Left:* Representative confocal acquisitions of growth cones of control neurons (CT, no Triton X100) and permeabilized (+Triton X100) Venus positive neurons (green), immunostained for endogenous HTT (grey) and microtubules (Tyrosinated tubulin, magenta). *Right:* Quantification of mean fluorescence intensity of indicated signals within growth cones after permeabilization. Unpaired t-test, **p <0.01; ***p <0.001 (Venus: 26.61 ±8.36; HTT: 143.50 ±56.04; Tyr-tub: 257.50 ±105.30; n=11 growth cones/condition). Mean ±SD are shown. (**B**) *Left:* Representative confocal images of growth cones of control neurons (CT, no Triton X100) and permeabilized (+Triton X100) Venus positive neurons (green), immunostained for endogenous HTT (grey) and F-actin (phalloidin, magenta). *Right:* Quantification of mean fluorescence intensity of indicated signals within growth cones after permeabilization. Unpaired t-test, *p <0.05, **p <0.01 (Venus: 24.88 ±9.33; HTT: 134.50 ±45.80; Phalloidin: 76.88 ±30.93; n=12 growth cones/condition). Mean ±SD are shown. (**C**) *Left:* Cartoon illustrating co-sedimentation assay at high speed of cortices lysate and microtubules. *Middle:* Representative western-blot of co-sedimentation assay from P0 cortical lysates immunoblotted for the presence of endogenous HTT and a-tub. *Right:* Quantification of the amount of HTT in absence or presence of microtubules (MTs). Paired t-test, ***p <0.001 (No MTs: 0.42 ±0.19; + MTs: 1.18 ±0.11; n=5 independent experiments). Mean ±SD are shown (**D**) *Left:* Cartoon illustrating co-sedimentation assay at high speed of cortices lysate and F-actin. *Middle:* Representative western-blot of co-sedimentation assay from P0 cortical lysates immunoblotted for the presence of endogenous HTT and a-actin. *Right:* Quantification of the amount of HTT in absence or presence of F-actin. Paired t-test, **p <0.01 (No F-actin: 0.37 ±0.22; + F-actin: 1.07 ±0.26; n=5 independent experiments). Mean ±SD are shown.

Thus, HTT associates with the actin and microtubule cytoskeleton but a larger proportion of HTT associated with F-actin than with microtubules.

### HTT binds directly to F-actin

We asked whether the purified human full-length (FL) wild-type HTT (23 glutamines) (Kim et al. 2021) interacts directly with F-actin. We incubated FL-HTT with and without F-actin and performed high-speed (100,000 × g) co-sedimentation (Fig. 3A). FL-HTT sedimented specifically in the pellet with F-actin, while it remained in the supernatant in the control without F-actin (Fig. 3B). We determined the strength of the interaction by analyzing the fraction of FL-HTT found in the pellet with increasing concentrations of F-actin (Fig. 3C). The calculated equilibrium dissociation constant (Kd_app_), which corresponds to the concentration of F-actin required to bind 50% of FL-HTT, was 357 ± 94 nM. Transmission electron microscopy showed FL-HTT molecules attached to actin filaments (Fig. 3D).

**Figure 3.**
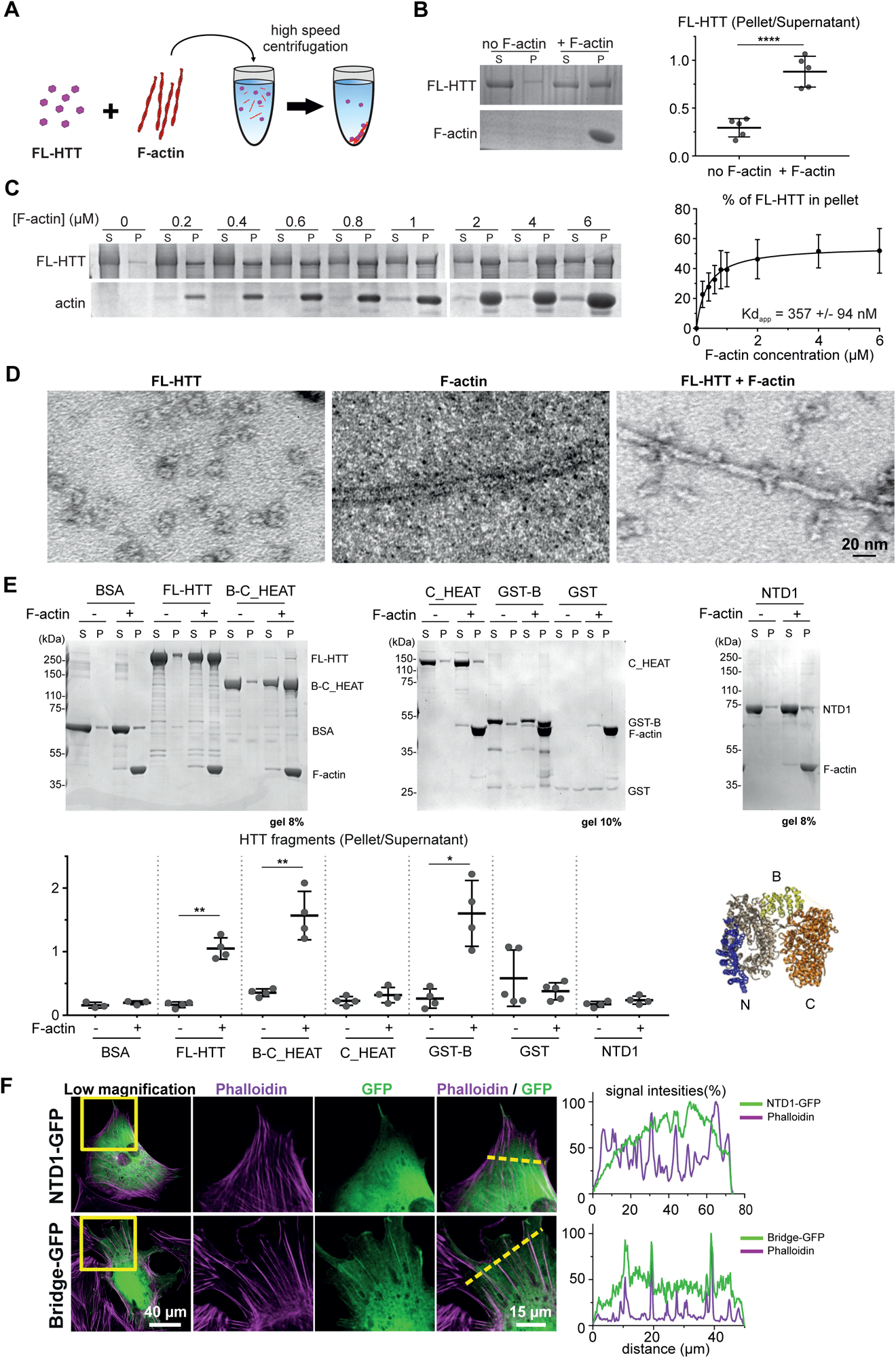
HTT binds actin filaments in a direct manner. (**A**) Cartoon illustrating co-sedimentation assay with purified FL-HTT (magenta) and F-actin (red). (**B**) *Left:* Representative SDS-PAGE stained with Coomassie blue. S, supernatant; P, pellet. *Right:* Graph shows the amount of FL-HTT found in the pellet with or without F-actin. **** p <0.0001, paired t-test (No F-actin: 0.29 *±*0.10; + F-actin: 0.88 *±*0.16; n=5 independent experiments). Data are presented as mean *±*SD. (**C**) Determination of FL-HTT binding affinity for F-actin. Representative SDS-PAGE of FL-HTT incubated with an increasing amount of F-actin. Graph to the right shows the bound fraction of FL-HTT expressed as the percent of total FL-HTT, Pellet+Supernatant relative to F-actin concentration. Kd_app_, calculated dissociation constant at equilibrium. n=5 independent experiments. Data are presented as mean *±*SD. (**D**) Representative electron micrograph of negative staining of FL-HTT, F-actin and F-actin incubated with FL-HTT. (**E**) *Up:* Representative SDS-PAGE stained with Coomassie blue of high-speed co-sedimentation assay with the different HTT fragments and F-actin. S, supernatant; P, pellet. *Bottom right:* Quantification of the amount of HTT fragments found in the pellet in presence or absence of F-actin. Pellet/Supernatant. Paired t-test, *p <0.05; **p <0.01 (BSA No F-actin: 0.16 ±0.05; BSA + F-actin: 0.20 ±0,03; FL-HTT No F-actin: 0.17 ±0.05; FL + F-actin: 1.05 ±0.17; B-C_HEAT No F-actin: 0.35 ±0.06; B-C_HEAT + F-actin: 1.57 ±0.38; C_HEAT No F-actin: 0.23±0.07; C_HEAT + F-actin: 0.32 ±0.12; GST-B No F-actin: 0.26 ±0.15; GST-B + F-actin: 1.60 ±0.52; GST No F-actin: 0.58 ± 0.44; GST + F-actin: 0.38 ±0.13; NTD1 No F-actin: 0.17 ±0.05; NTD1 + F-actin: 0.24 ±0.06; at least n=3 independent experiments). Mean ±SD are shown. *Bottom left:* FL-HTT structure deposited online and available in the Protein Data Bank archive (PDB: E6Z8). Structure of HAP40 was hidden and NTD1, Bridge and C_HEAT domains of HTT were colored in blue, yellow, and orange, respectively. (**F**) *Left:* Representative confocal images of MEFs transfected with NTD1-GFP or Bridge-GFP and immunostained with phalloidin to label F-actin. Yellow boxes show the area of higher magnification and yellow dotted lines indicate the axis along which the representative line-scan analyses were performed. *Right:* Representative line-scan analyses (relative fluorescence intensity) of indicated immunostainings.

To decipher how HTT interacts with F-actin, we used recombinant proteins corresponding to different HTT deletion mutants. HTT contains three large domains: N_HEAT (1-1,684 a.a.), C_HEAT (2,091-3,144 a.a.), and the Bridge domain linking the HEAT domains (Vijayvargia et al. 2016; Guo et al. 2018). We produced one HTT fragment containing the Bridge to C_HEAT domains (B-C_HEAT; 1887 to 3144 a.a.); a second containing just the Bridge domain tagged with GST (GST-B: 1887-2098 a.a.); and a third containing the C_HEAT domain (C_HEAT; 2098-3144 a.a.) (Fig. 3E, Supplementary Fig. 2). We were not able to isolate the entire N_HEAT domain, but only a fragment (1-606 a.a.), which was previously defined as N-terminal domain 1 (NTD1) using crosslinking mass spectrometry (MS) (Vijayvargia et al. 2016). We assessed the ability of each of these fragments to bind to F-actin by high speed cosedimentation assays (Fig. 3E). When HTT fragments were incubated with F-actin, we observed that only the B-C_HEAT and GST-B fragments co-sedimented with F-actin, as did FL-HTT. We confirmed this result by using a glycerol gradient ultracentrifugation (Supplementary Fig. 2). This procedure allows free HTT fragments to remain in fractions of low glycerol concentrations, while the F-actin and the filament-bound HTT enter fractions of high glycerol concentrations (Supplementary Fig. 2). When HTT fragments were incubated with F-actin, we observed that only the B-C_HEAT and GST-B fragments co-migrated with F-actin in high glycerol concentration fractions (as did FL-HTT).

Next, we produced plasmids encoding for the fragments NTD1 and Bridge tagged with GFP fragments, named NTD1-GFP and Bridge-GFP, respectively. We transfected Mouse Embryonic Fibroblasts (MEFs) with these constructs and performed immunostaining against GFP and labeled F-actin using phalloidin (Fig. 3F). Confocal microscopy revealed that NTD1-GFP was mostly diffuse in the cytoplasm while the Bridge-GFP staining adopted a filamentous pattern that colocalized with phalloidin. Line scan analyses show that the mean fluorescence intensity of NTD1-GFP homogeneously distribute throughout the cell without correlation with the one of phalloidin, while distinct pikes could be observed for Bridge-GFP that were concomitant with the ones of phalloidin (Fig. 3F).

Together, these results unequivocally show that HTT is an actin-binding protein and uses its Bridge domain to interact with actin filaments.

### HTT oligomers bundle actin filaments

We next investigated whether HTT regulates F-actin dynamics. We used Total Internal Reflection Fluorescence (TIRF) microscopy to visualize the polymerization of actin covalently bound to the ATTO565 fluorescent group (ATTO565-actin) (Fig. 4A). TIRF microscopy experiments revealed that the presence of FL-HTT enabled the polymerization of unbranched filaments and that HTT induces the formation of bundles, visualized as large objects with high fluorescence intensity (Fig. 4A). Quantifications show that higher concentrations of FL-HTT induced a greater proportion of the F-actin network to organize into bundles (Fig. 4B). Measuring the mean fluorescence intensity of the ten brightest objects—single filaments in the control conditions and bundles for 125 nM and 500 nM FL-HTT for each time point—showed that bundles induced by 500 nM FL-HTT had a greater mean fluorescence intensity than those induced by 125 nM FL-HTT (Fig. 4B). This suggests that the number of filaments integrated into bundles increases with the amount of HTT present. Consistently, confocal microscopy observations of pre-polymerized F-actin stabilized with the fluorescent Phalloidin-ATTO488 after 1h incubation with high concentration (750 nM) of FL-HTT revealed the formation of large F-actin bundles containing numerous filaments assembled in 3 dimensions (Supplementary Fig. 3A).

**Figure 4.**
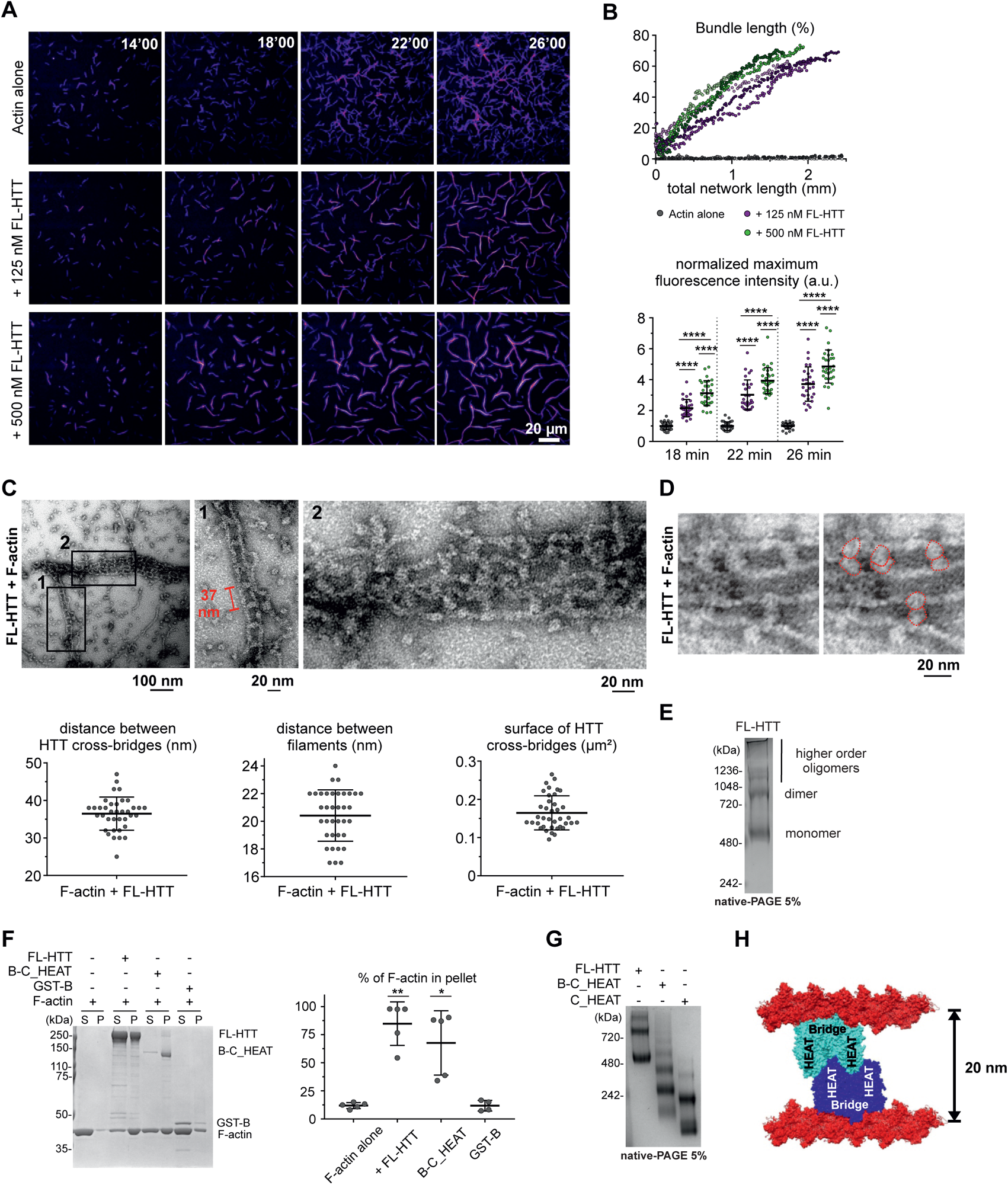
HTT oligomerizes to bundle actin filaments. (**A**) Representative TIRF time-lapse images of ATTO565-actin polymerization with different FL-HTT concentrations. (**B**) *Up:* Graph shows the fraction of F-actin in bundles relative to the total actin network length. Three representative curves per condition are displayed in shades of green, magenta and grey. *Bottom:* Graph shows the mean fluorescence intensity of the 10 brightest filaments/bundles at several time points in the absence or presence of different FL-HTT concentrations; **** p <0.0001, One-way ANOVA followed by Tukey’s multiple comparisons post hoc test (for CT: 1 at all times; for 125 nM: 2.155; 3.019; 3.712; for 500 nM: 3.124; 3.933; 4.850; n=3 replicates with at least 10 filaments per condition per replicate). Data are presented as mean *±*SD. (**C**) Representative electron micrograph of negative staining showing F-actin bundles covered by FL-HTT molecules. Magnification of region *1* shows the regular spacing between FL-HTT cross-bridges. Magnification of region *2* shows the regular spacing between filaments within a bundle. Graphs below the images show the spacing between HTT cross-bridges (36.5 *±*4.4), the distance between actin filaments within a bundle (20.4 *±*1.9), and the surface covered by HTT cross-bridges (164.7 *±*44.6). (**D**) Representative electron micrograph of negative staining showing FL-HTT cross-bridges formed by pairs of globular structures. *Left:* original acquisition. *Right:* addition of dotted lines delimitating pairs of spherical objects. (**E**) Representative Native-PAGE showing oligomer of FL-HTT. (**F**) Representative SDS-PAGE of low-speed co-sedimentation assay of F-actin in the presence of different HTT fragments. S, supernatant; P, Pellet. Graph below shows the amount of F-actin found in the pellet relative to the different HTT fragments; **p <0.01, *p <0.05, One-way ANOVA followed by Tukey’s multiple comparisons post hoc test (F-actin alone: 11.9 *±*2.7; FL-HTT: 84.8 0 *±*19.4; B-C_HEAT: 67.6 *±*28.6; GST-B: 11.8 *±*4.8; at least n=4 independent experiments). Data shown as mean *±*SD. (**G**) Representative Native-PAGE showing oligomers of FL-HTT, B-C_HEAT and C_HEAT. (**H**) Cartoon illustrating the model proposed for HTT-dependent bundling of F-actin. HTT binds a single actin filament through its bridge domain and needs to dimerize through either its N_ or C_HEAT domains to crosslink a second actin filament.

To confirm the HTT bundling activity biochemically, we incubated pre-polymerized F-actin with FL-HTT and performed low-speed (15,000 x g) centrifugation to pellet the bundles while leaving single filaments in the supernatant (Elie et al. 2015). We then measured the fraction of F-actin found in the pellet. In control samples containing F-actin alone (single filaments), a minimal amount of filaments were found in the pellet after low-speed centrifugation. However, F-actin was gradually enriched in the pellet with increasing doses of FL-HTT, with 750 nM of HTT leading to 68% of the initial 2 µM of F-actin being bundled (Supplementary Fig. 3B).

We used transmission electron microscopy to analyze at higher resolution the bundles formed by FL-HTT after 1h incubation with pre-polymerized F-actin (Fig. 4C). We first observed that adjacent filaments were linked by FL-HTT cross-bridges spaced 37 nm apart, a distance that corresponds to half a pitch of an F-actin helix (Holmes et al. 1990). The mean spacing between two adjacent actin filaments within a bundle was 20 nm (Fig. 4C). FL-HTT cross-bridges covered a surface of approximately 164 nm² whereas the theoretical surface covered by a single FL-HTT molecule, estimated by an ellipse of 6 x 5 nm radius (Jung et al. 2020), is approximately ∼94.2 nm². This discrepancy suggested to us that HTT might oligomerize to exert its bundling function. In fact, the FL-HTT cross-bridges were made of two globular parts that could correspond to dimers (Fig. 4D, red circles). Native PAGE showed that FL-HTT migrated at molecular weights corresponding to monomers, dimers and higher-order oligomers (Fig. 4E). We also observed dimers of endogenous HTT from extracts of P0 developing cortices, as previously described (Li et al. 2006) (Supplementary Fig. 3C). We detected the presence of homodimers from concentrations as low as 50 nM of FL-HTT, which is lower than the concentrations used to visualize bundles in our previous electron microscopy, TIRF microscopy and bundling assays (Supplementary Fig. 3D). These results suggested that HTT might oligomerize to exert its bundling function.

As GST-B and B-C_HEAT fragments can interact with F-actin as FL-HTT does (Fig. 4), we next asked whether they also retained the ability to crosslink actin filaments into bundles (Elie et al. 2015) (Fig. 4F). Low-speed centrifugation revealed F-actin in the pellet with FL-HTT and the B-C_HEAT fragment but not with the GST-B fragment. Native-PAGE showed that the B-C_HEAT fragment was as capable as FL-HTT of forming dimers and that the C_HEAT domain was sufficient to do so (Fig. 4G). As the NTD1 fragment was also able to oligomerize, the oligomers obtained with the B-C_HEAT fragment could be of a different nature from those formed by FL-HTT (Supplementary Fig. 3E).

These results suggest a model for HTT-dependent bundling of F-actin in which HTT uses its Bridge domain to interact with a single actin filament while it dimerizes through its N_HEAT and/or C_HEAT domains to crosslink adjacent filaments into bundle with 20 nm inter-filament spacing (Fig. 4H).

### HTT assembles a specific set of proteins on F-actin

We next wondered which physiological function the HTT/F-actin partnership may serve. As HTT responds to neural physiology by scaffolding protein complexes on vesicles with great specificity (Saudou and Humbert 2016), we hypothesized that it could serve a similar function on F-actin. This would accord with an emerging view in the actin field that the actin cytoskeleton adopts different identities in different subcellular structures by adopting different conformations and associating with specific proteins in each context (Boiero Sanders et al. 2020). We therefore performed mass spectrometry to compare the proteins that co-sediment with F-actin in the absence or presence of HTT (Fig. 5A). We incubated F-actin with or without FL-HTT *in vitro*. We next added a protein extract from DIV4 cortical neurons previously depleted of endogenous HTT to make sure that the endogenous HTT did not interfere with our study. We first verified that Hdh^loxlox^ neurons infected with a lentivirus expressing the CRE-recombinase (LV-CRE) showed a marked reduction of HTT protein level (Fig. 5B). We then verified that the purified actin cytoskeleton was intact after incubation with lysate, with or without FL-HTT, and observed numerous filaments after co-sedimentation (Fig. 5C). FL-HTT remained bound to F-actin after the lysate addition, as reflected by the large quantity of FL-HTT found in the pellet with F-actin after co-sedimentation, relative to the small amount of FL-HTT found in the pellet without F-actin (Fig. 5D). We prepared samples containing only F-actin and others containing both F-actin and FL-HTT (Fig. 5A), using a lysate-only control to assess nonspecific proteins in the pellet fraction after ultracentrifugation and send them for proteomic-based mass-spectrometry analysis (Fig. 5E).

**Figure 5.**
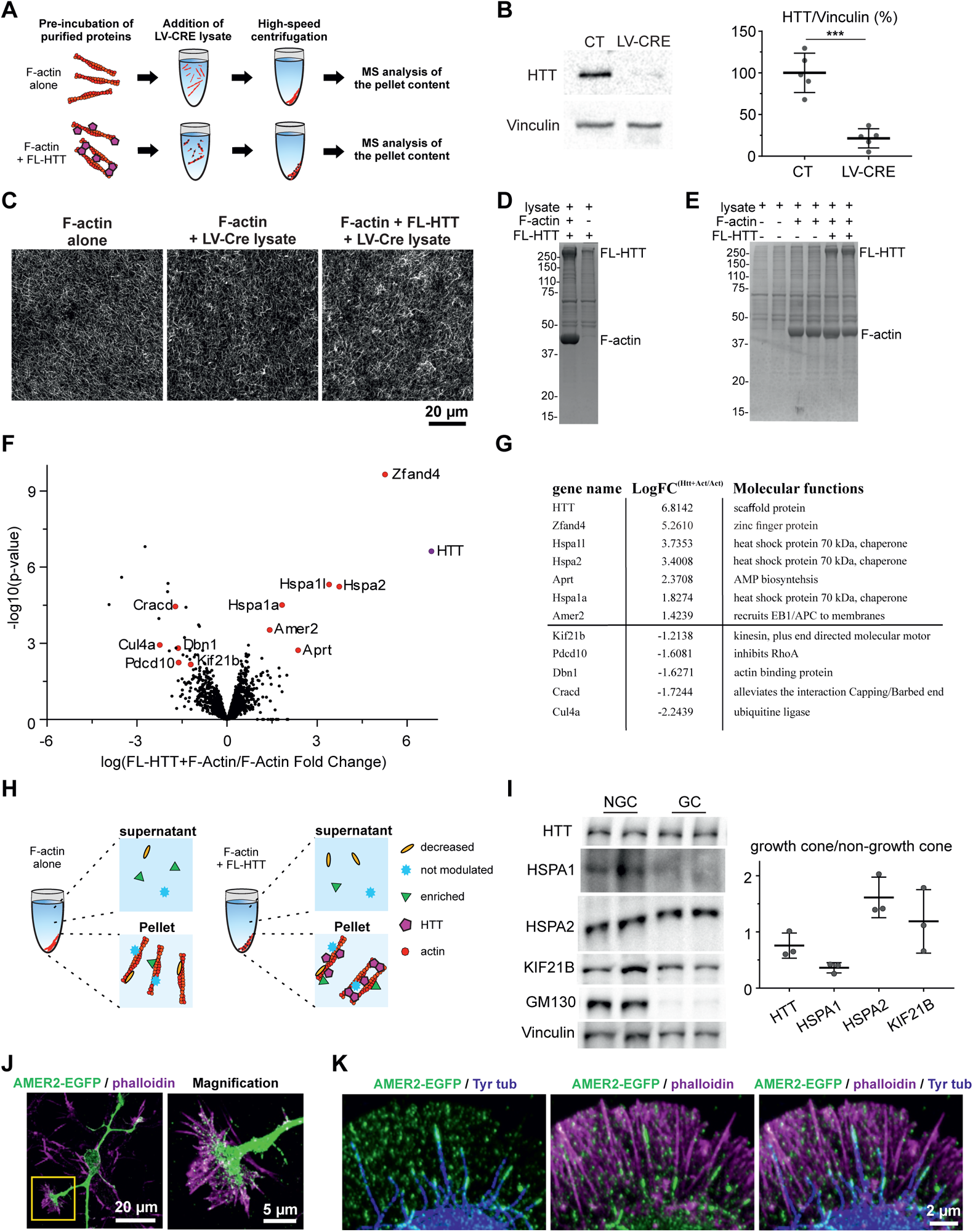
HTT establishes a specific protein network on F-actin. (**A**) Cartoon illustrating the experimental procedure for mass spectrometry analysis. (**B**) Representative western blot of lysates from DIV4 Hdh^lox/lox^ neurons infected with LV-GFP (CT) or LV-CRE (LV-CRE). Graph shows the relative quantities of HTT and Vinculin. ***p <0.001, paired t-test (CT: 100.0 ±23.6; CRE: 21.5 ±11.5; n=5 independent experiments). Data are presented as mean ±SD. (**C**) Representative confocal images of pre-polymerized F-actin with or without FL-HTT after 15 min incubation in lysate. (**D**) Representative SDS-PAGE stained with Coomassie blue of pellet fractions obtained after high-speed centrifugation of lysate incubated with FL-HTT or FL-HTT/F-actin. (**E**) Representative SDS-PAGE stained with Coomassie blue showing the pattern of proteins obtained in the pellet after high-speed centrifugation of the lysate alone, F-actin+lysate, and HTT+F-actin+lysate. (**F**) Volcano plot representing the differential proteins associated with F-actin/FL-HTT or F-actin alone. Red dots indicate proteins significantly modulated (fold change ≥ 2 and limma p-value ≤ 0.01). (**G**) The dozen proteins most significantly modulated between F-actin and F-actin/FL-HTT network. (**H**) Cartoon illustrating that the presence of FL-HTT on F-actin allows specific proteins to be pulled down. (**I**) *Left*: Representative western blot of growth cone (GC) and non-growth cone fractions (NGC) obtained with growth cone fractionation protocol. *Right*: Quantification of protein abundance in growth cone fraction relative to non-growth cone fraction after normalization of protein levels on Vinculin. (HTT: 0.76 ±0.22; HSPA1: 0.36 ± 0.09; HSPA2: 1.61 ±0.36; KIF21B: 1.19 ±0.57; n=3 fractionations). Data are presented as Mean ±SD. (**J**) Confocal image of a neuron expressing AMER2-EGFP (green) and immunostained for F-actin (phalloidin, magenta). (**K**) Representative airyscan confocal image of the peripheral domain of a growth cone of a neuron expressing AMER2-EGFP (green) and immunostained for F-actin (phalloidin, magenta) and microtubules (Tyr tub, blue).

After high-speed ultracentrifugation, we found 120 proteins to be significantly more abundant in the presence of F-actin than in the lysate-only pellet (fold-change ≥ 2 and limma p-value ≤ 0.01; Supplementary Fig. 4A). As expected, among these were numerous known actin-binding proteins, and Gene Ontology (GO) analysis revealed a significant enrichment for terms related to the actin cytoskeleton (Supplementary Fig. 4A, Supplementary Table 1). Similarly, the 93 proteins that were significantly enriched in the pellet of the sample containing both HTT and F-actin also were enriched for actin-related proteins as compared to the lysate-only sample (Supplementary Fig. 4B, Supplementary Table 1). Fifty-nine proteins were common to both the F-actin and the HTT/F-actin proteomes, with no specific enrichment in GO terms between the two.

Most notably, we found 12 proteins for which the abundance in the F-actin pellets was significantly modulated by the presence of HTT on actin: seven of them were enriched, and five were less abundant in the presence of HTT (Fig. 5F-H). Several of them are known to act at the level of the cytoskeleton: The Heat Shock Protein members HSPA1 and HSPA2 localize to F-actin in the cytoplasm of A6 mesoangioblasts cells and *Dictyostelium discoideum* (Xiang and Rensing 1999; Turturici et al. 2008) and cross-link actin filament tips; PDCD10 (Programmed Cell Death 10), also known as CCM3 (Cerebral Cavernous Malformation 3) localizes at cell adhesion sites and inhibits RhoA (Louvi et al. 2014; Valentino et al. 2021); CRACD (Capping Protein Inhibiting Regulator Of Actin Dynamics) sequesters Capping protein from actin filament tips (Jung et al. 2018); the kinesin motor KIF21B limits microtubule growth (Hooikaas et al. 2020); AMER2 (APC membrane recruitment protein 2) anchors microtubule plus-ends under the plasma membrane (Pfister et al. 2012; Siesser et al. 2012); Drebrin1, an actin bundling protein, crosslinks F-actin with microtubules ends (Geraldo et al., 2008; Worth et al., 2013). Several of these candidates could function in growth cones. Loss of the kinesin motor KIF21B (Asselin et al. 2020) and of Drebrin1 leads to axonal growth defects (Geraldo and Gordon-Weeks 2009). AMER2 binds microtubule plus ends through its interaction with EB1 (End-Binding 1) and APC (Adenomatous Polyposis Coli), both proteins already described in growth cones (Mimori-Kiyosue et al. 2000; Koester et al. 2007; Purro et al. 2008).

We western blotted the protein contents of growth cones from extracts of developing cortices isolated by sucrose gradient fractionation (Fig. 5I). HSPA1 was more abundant in non-growth cone fractions, but KIF21B and HSPA2 were present in both non-growth cone and growth cone fractions, supporting the notion that they play a role in growth cones. Because there is no antibody available to detect AMER2, we expressed a plasmid encoding AMER2 tagged with EGFP (AMER2-EGFP) in cortical neurons and traced AMER2-EGFP through the cell and growth cones (Fig. 5J). AMER2-EGFP was visible in comet-like streaks near the microtubule plus ends, as previously described (Pfister et al. 2012; Siesser et al. 2012), at the interface between exploratory microtubules and actin filaments (Fig. 5K).

Altogether, HTT appears to modulate the association of specific proteins with F-actin in growth cones as well as in other subcellular compartments.

### HTT depletion impairs axonal growth

To address the functional consequences of our findings, we depleted HTT by electroporating DIV0 primary cortical neurons from Hdh^lox/lox^ embryos with a plasmid encoding either the reporter gene Td-tomato or Td-tomato plus the CRE-recombinase. We labeled F-actin with phalloidin, used antibody directed against tyrosinated-tubulin to visualize microtubules and measured several parameters: growth cone areas, actin crown perimeter, the number of filopodia and the length of exploratory microtubules (Fig. 6A). Overall, loss of HTT seems to loosen the architecture of the growth cone: DIV4 HTT-depleted neurons displayed larger growth cone areas with fewer filopodia (Fig. 6B). The HTT-depleted neurons also had longer exploratory microtubules.

**Figure 6.**
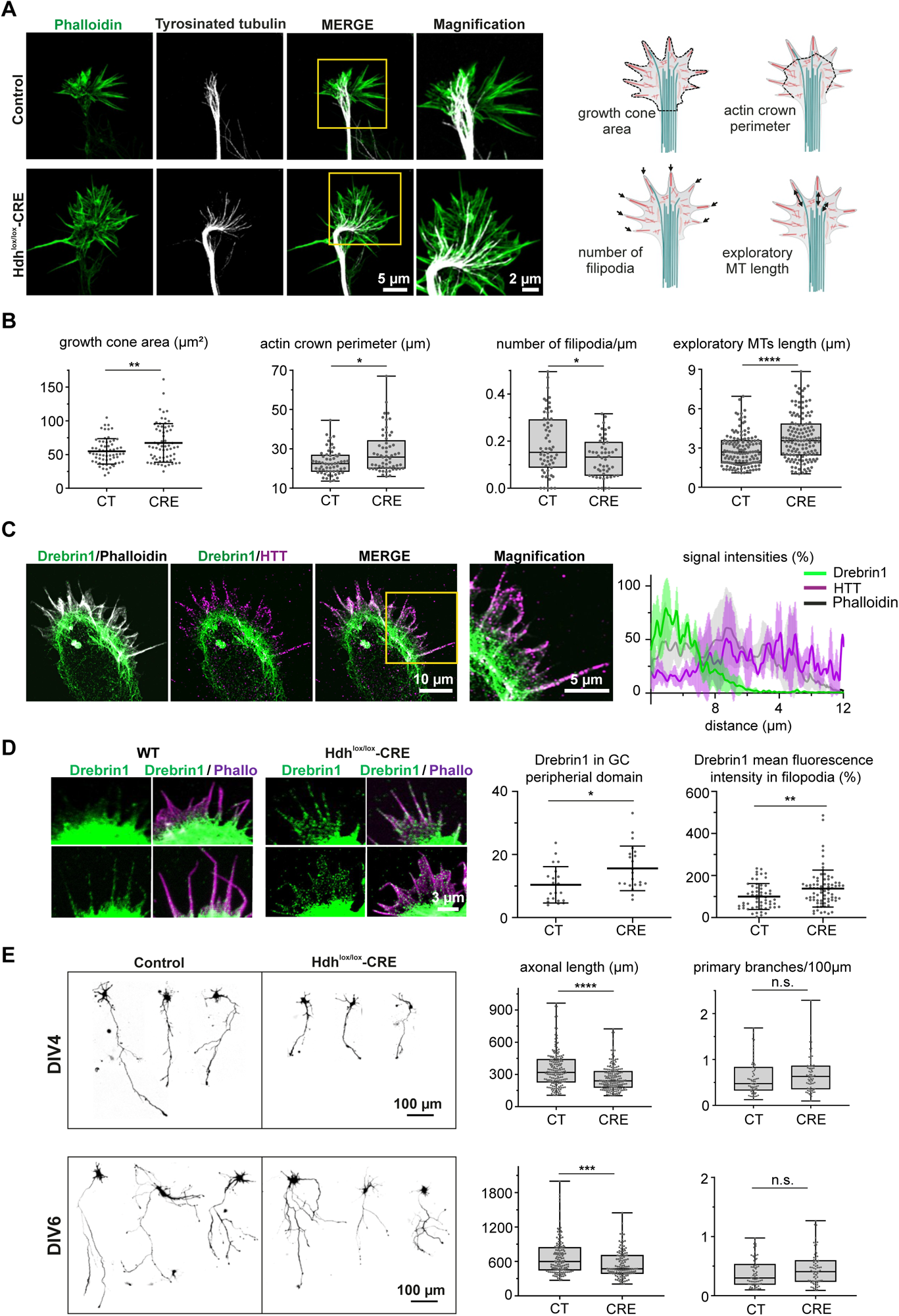
HTT is necessary for cytoskeleton organization in growth cones. (**A**) Representative airyscan confocal images of DIV4 CT and CRE growth cones labeled for F-actin (phalloidin, green) and Tyrosinated tubulin (grey). Drawings illustrate the parameters measured. (**B**) Growth cone area, **p <0.01, unpaired t-test (CT: 54.8±18.9 CRE: 67.3 ±28.6; at least 63 growth cones/condition). Actin crown perimeter, *p <0.05, Mann-Whitney test (CT: 22.5, CRE: 25.8; at least 57 growth cones per condition); density of filopodia, p <0.05, Mann-Whitney test (CT: 0.15; CRE: 0.13; at least 52 growth cones per condition); length of exploratory microtubules, ****p <0.0001, Mann-Whitney test (CT: 2.69 CRE: 3.56 at least 42 growth cones/condition). n=6 independent experiments with at least 7 growth cones/genotype. Normally and non-normally distributed data are presented as mean ±SD and in box-whisker plots, respectively. (**C**) *Left:* Representative airyscan confocal image of a DIV4 growth cone immunostained for endogenous HTT (magenta), endogenous Drebrin1(green) and F-actin using phalloidin (grey); the yellow box shows the magnified area. *Right:* Representative line-scan analysis (relative fluorescence intensity) of indicated immunostainings across the transitional domain until the end of the peripheral domain of growth cones; n=4 growth cones. (**D**) *Left:* Representative confocal images of DIV4 CT and CRE growth cones immunostained for endogenous Drebrin1 (green) and F-actin using phalloidin (magenta). *Right:* Quantifications of Drebrin1 mean fluorescence intensity of CT and CRE measured in the peripheral domain, *p <0.05, unpaired t-test (CT:, 0710.39±5.78 CRE: 15.60 ±xx.x; at least 5 growth cones/condition/experiment); or in filopodia, **p <0.01, unpaired t-test (CT: 100.00±61.65 CRE: 137.80 ±87.94; at least xx filopodia/condition/experiment). n=3 independent experiments. Data are presented as Mean ±SD. (**E**) Representative confocal images of CT and CRE neurons. The graphs quantify axonal length and branching. Length at DIV4, ****p <0.0001, Mann-Whitney test (CT: 319.3; CRE: 240.1; n=3 independent experiments with at least 40 axons per condition and experiment). Branching at DIV4, not significant (n.s.), Mann-Whitney test (CT: 0.48; CRE: 0.63; n=3 independent experiments with at least 20 axons per condition and experiment). Length at DIV6, ***p <0.001, Mann-Whitney test (CT: 599.10; CRE: 470.70; n=3 independent experiments with at least 40 axons per condition and experiment). Branching at DIV6 n.s., Mann-Whitney test (CT: 0.30; CRE: 0.41; n=3 independent experiments with at least 20 axons per condition and experiment). Data are presented as box-whisker plots.

We were particularly interested in Drebrin1, which is decreased on F-actin in presence of HTT (Fig. 5F-H) but was shown to interact with F-actin in growth cones (Geraldo and Gordon-Weeks 2009; Worth et al. 2013). We used airyscan confocal microscopy to investigate the respective localization of endogenous HTT, Drebrin1 and actin in growth cones of DIV4 cortical neurons (Fig. 6C). While HTT mostly localized in the peripheral domain on filopodia, we observed that Drebrin1 is strongly enriched at the level of the growth cone transitional domain on actin arcs, as reported previously (Geraldo and Gordon-Weeks 2009). This was confirmed by line scan analyses across the transitional domain until the end of the peripheral domain of growth cones, showing that Drebrin1 fluorescence signal is mostly contained within the transitional domain and that HTT fluorescence signal extend beyond and reaches its maximal intensity within the peripheral domain (Fig. 6C). Thus, HTT and Drebrin1 segregate on different F-actin structures in the peripheral and transitional domains of growth cones, on actin bundles and contractile acto-myosin arcs, respectively.

We next analyzed the consequences of HTT depletion on Drebrin1 localization in growth cones. We compared Drebrin1 localization in growth cones of Hdh^loxlox^ neurons infected with LV-CRE compared to non-infected Hdh^loxlox^ neurons (Fig. 6D). We performed immunostaining for endogenous Drebrin1 and labeled F-actin using phalloidin and measured Drebrin1 mean fluorescence intensity in the peripheral domain of growth cones and within filopodia only (Fig. 6D). Quantifications revealed an increased mean fluorescence intensity of Drebrin1 staining in both the full peripheral domain and filopodia of HTT-depleted growth cones. These analyses show that HTT and Drebrin1 localize on different F-actin structures in growth cones and that HTT is necessary for the proper localization of Drebrin1, segregating it on actin arcs.

Given the evident cytoskeletal defect and mislocalization of Drebrin1, we examined axonal growth which relies on the integrity of the growth cones. Neurons electroporated with the CRE-recombinase showed significantly shorter axons at DIV4 and 6, although the number of primary branches was similar (Fig. 6E).

We conclude that loss of HTT impairs the cytoskeletal organization in the growth cone with functional consequences on axonal growth.

## Discussion

Understanding the functions of HTT revealed how it could scaffold large multi-protein complexes, implying HTT in a variety of biological processes (Saudou and Humbert 2016). Here, we report a direct function for HTT that acts as an actin-binding protein, conferring that cytoskeletal component a specific structure and proteomic identity. Together with the recent discovery that HTT participates in the post-translational modification of actin with its partners HIP1 and SETD2 (SET domain containing 2) (Seervai et al. 2020), the current study lends weight to recent developments in the actin field indicating that different populations of actin filaments have distinct identities (Boiero Sanders et al. 2020). Thus, HTT role on the actin cytoskeleton could contribute to the regulation of the global cellular integrity by regulating the cell physiology from its shape and structure, to signaling and transport.

Previous studies indicated that HTT is involved with actin (Lynch et al. 2007; Munsie et al. 2011; Miller et al. 2012; Tourette et al. 2014; Tousley et al. 2019a). As suggestive as all these studies have been, evidence of a direct role for HTT in organizing the cytoskeleton has been lacking. We employed purified recombinant HTT in its full length and deletion mutants to identify that HTT interacts with F-actin through its Bridge domain and oligomerize to bundle actin filaments. While most of the interactions with HTT partners have been reported to occur via its N-HEAT (Saudou and Humbert 2016), we provide deeper insights in our understanding of HTT biology at the molecular scale and one physiological function for HTT dimers (Li et al. 2006).

We identify the global proteome associated with HTT/F-actin complexes and show that HTT modulates the association of several proteins with F-actin. Several mechanisms could intervene here. Upon HTT association with F-actin, the enrichment of proteins could result from an increased accessibility to their binding sites on the filament or from a highly context-specific interaction with HTT scaffolding actin. As important are the proteins whose association with F-actin were limited by HTT. There may be competition with favored proteins and HTT for a common binding site on the filament or a change in accessibility to the filament because of a conformational change of the cytoskeleton itself (Winkelman et al. 2016; Boiero Sanders et al. 2020; Harris et al. 2020). For example, Drebrin1 binding could be disfavored because HTT cross-bridges were regularly spaced at 37 nm, suggesting that the pitch of the F-actin helix is unchanged, and Drebrin1 prefers a longer pitch (Sharma et al. 2012). Another possibility is that the differential inter-filament spacing within bundles formed by HTT or Drebrin1, of 20 nm or 12 nm (Worth et al. 2013) respectively, influences protein sorting as observed between the large and compact bundlers α-actinin and Fascin (Winkelman et al. 2016).

Local specification of the proteome could conceivably be required to guide growth cone navigation, a process that relies on asymmetric remodeling of the cytoskeleton and for which the coupling of the exploratory microtubules with F-actin is crucial. For example, both Drebrin1 and AMER2 bind microtubule ends through interaction with EB proteins (Geraldo and Gordon-Weeks 2009; Pfister et al. 2012; Siesser et al. 2012; Worth et al. 2013) but HTT favors the attachment of AMER2 on F-actin while antagonizing Drebrin1. HTT might help regulate the coupling of microtubules to actin by controlling the association of specific F-actin/microtubule cross-linkers on actin filaments.

As our proteomic-based mass-spectrometry approach was performed on total cell extract, it gave an overview of the protein networks on the actin cytoskeleton in which HTT can be involved that could be extended to other actin structures in other spatial or temporal contexts. For instance, HSP70 may exert a protective role for actin filaments, as HSP70 relocalizes to F-actin shortly after heatshock (Xiang and Rensing 1999; Turturici et al. 2008). HTT absence and mutation impairs the formation and persistence of nuclear ADF/Cofilin actin rods forming upon heatshock (Munsie et al. 2011). Thus, HTT could participate to the stress response stress by favoring the presence of HSP70 on the actin cytoskeleton in several cell compartments. Consistent with a role on other actin structures, HTT has been observed on anti-parallel contractile acto-myosin structures found in stress fibers and rings (Tousley et al. 2019a) (Tousley 2019; Atwal 2011) that are necessary for cell migration and cytokinesis, respectively (Svitkina 2018; Zaidel-Bar 2015). Interestingly, HTT depletion triggers to the mislocalisation of its interactor α-actinin, an actin bundling protein localizing at stress fibers, and leads to the failure of Myosin-II motors to localize at the acto-myosin rings (Tousley et al. 2019a). In agreement with a defective contractility of stress fibers, analyses of neuruloids revealed that HTT depletion and mutation trigger morphologic alterations that are recapitulated by a drug inhibiting Myosin II (Haremaki et al. 2019). Altogether, these data set HTT as a key regulator of the actin cytoskeleton.

Proteomic analyses of post-mortem HD brains revealed a general dysregulation of actin regulatory pathways (Ratovitski et al. 2012; Tourette et al. 2014; Haremaki et al. 2019). Together with our recent work showing that microtubules in HD growth cones are disorganized (Capizzi et al. 2022), the current study hints that derangement of the cytoskeleton accompanies alterations in intracellular transport and other HTT-related functions in HD. Numerous developmental and neurophysiological processes rely on the close cooperation of these two cytoskeleton components, from interkinetic nuclear migration, cell division, and radial migration to axonal growth and tissue maintenance, all of which are altered in HD (Godin et al. 2010; Molina-Calavita et al. 2014; Saudou and Humbert 2016; Barnat et al. 2017; Barnat et al. 2020). Disorganization of the cytoskeleton itself could thus underlie numerous pathogenic alterations observed in HD. This study supports the hypothesis that HD is neurodegenerative disease with a developmental component which may originate from cytoskeletal alterations (Lasser et al. 2018; Munoz-Lasso et al. 2020).

This study provides a first glimpse of HTT interactions with the actin cytoskeleton. Future investigations with other actin structures would be required to address the precise role of HTT and the different candidates for which the association with F-actin is modulated by HTT and to understand to what extent the pathogenic HTT mutation alters these processes.

## Materials and Methods

### Primary culture

Cortices were dissected from E15.5-E16.5 embryos and digested in Papain enzyme solution. Papain was inactivated using 10% FBS and cells were washed with opti-MEM-glucose. Cells were cultured on poly-L-lysine matrices in Neurobasal supplemented with 2% B27, 1% glutamax, 1% Pennicillin/Streptamycin and 10% FBS for 1 h and medium was replaced by Neurobasal containing 2% B27, 1% glutamax and 1% Pennicillin/Streptamycin. Neurons were incubated in 5% CO_2_ in a humidified incubator at 37°C. For western blot analysis, cells were plated at 2,000,000 cells in a P10 Petri dish and were lysed at indicated DIV.

For analysis of endogenous HTT expression and localization, we used embryos from the Swiss/CD1 mouse strain (Janvier Lab). For permeabilization assays, Swiss/CD1 neurons were electroporated/nucleofected at DIV0 with 3 µg of plasmid encoding for mVENUS (pCAG-mVENUS pSCV2, provided by J. Courchet Neuromyogene, Lyon, France) and mixed 1:1 with non-electroporated cells to favor cell survival, or with AMER2-EGFP (Provided by M.Sachs and J. Behrens, Nikolaus-Fiebiger-Zentrum, Erlangen, Germany) to assess AMER2 subcellular localization. To assess the effects of HTT depletion, Hdh^lox/lox^ mouse (Dragatsis and Zeitlin 2001) embryos were electroporated at DIV0 just before plating with 3 µg of the plasmid control encoding for Td-tomato alone (pCAG td Tomato, Addgene) or 3 µg of plasmid encoding for Cre-recombinase and reporter gene Td-tomato (pCAG td Tomato-Ires-Cre, Gage’s lab). To assess HTT downregulation by immunoblotting and analyze growth cone morphology, electroporated cells were mixed (1:1) with non-electroporated cells. For axonal length and branching analysis, electroporated cells were mixed (1:4) with non-electroporated cells. For immunofluorescence analysis, cells were plated at 800,000 cells/well of a 6-well plate and were fixed at DIV4.

### Western blot

Cultured cortical neurons were lysed at indicated DIV in RIPA buffer (20 mM Tris-HCl, 150 mM NaCl, 2.5 mM Sodium pyrophosphate, 1 mM EDTA, 1 mM EGTA, 1% Deoxycholate, 1% NP40) supplemented with protease inhibitors and phosphatase inhibitors (Sigma-Aldrich). Samples were centrifuged for 10 min at 10,000 times gravity (× g) at 4 °C insoluble materials and total protein concentration was measured by the Pierce^TM^ BCA protein assay (Thermo Fisher Scientific). 15-20 µg of samples was prepared in Laemmli buffer and run in 8% SDS-PAGE before being transferred onto nitrocellulose membrane at 320 mA for 2.5 h. After 1 h of blocking with Tris-buffered saline with 0.1% Tween detergent (TBST) and 5% BSA, membranes were incubated overnight with primary antibodies mouse anti-vinculin (Sigma-Aldrich) diluted 1:1000; mouse anti-Tau (Abcam) diluted 1:500; rabbit anti-α-skeletal-actin (Sigma-Aldrich) diluted 1:2000; mouse anti-a-tubulin (Sigma-Aldrich) diluted 1:2000; mouse anti-HTT (4C8, IGBMC, Strasbourg) diluted 1:500; mouse anti-JIP1 (B-7, Santa Cruz Biotechnology) diluted 1:500; mouse anti-GM130 (BD Transduction Laboratories) diluted 1:250; rabbit anti-KIF21B (Sigma-Aldrich) diluted 1:500; mouse anti-HSPA1A (StressGen Biotechnologies Corp.) diluted 1:500; rabbit anti-HSPA2 (Sigma-Aldrich) diluted 1:500. Washes were performed with TBST. Secondary antibodies Goat anti-mouse IgG1-HRP conjugate (SouthernBiotech) diluted 1:2500; anti-rabbit IgG H&L-HRP (Interchim) diluted 1:2500, wre incubated for 1 h at RT and membranes were revealed by using ECL reagents with the ChemiDoc imager (Bio-Rad Laboratories).

### Immunofluorescence

Cells were fixed at DIV4 with 0.5 % glutaraldehyde (Sigma-Aldrich), 0.1 % Triton-X100 (Sigma-Aldrich) in cytoskeleton buffer (10 mM MES, 138 mM KCl, 3 mM MgCl_2_, 2 mM EGTA, pH 6.1) supplemented with sucrose (10 %) and pre-warmed at 37 °C. Glutaraldehyde auto-fluorescence was quenched by 10 min incubation in sodium borohydride 1 mg/mL. Blocking was performed with PBS-5 % BSA for 1 h at RT. Primary antibodies were incubated overnight at 4 °C. Secondary fluorochrome-conjugated antibodies were applied for 1.5 h at RT and phalloidin (1:1000) for 30 min at RT. Glutaraldehyde autofluorescence was quenched by 10 min incubation with PBS supplemented with sodium borohydride (Sigma-Aldrich). Washes were done in PBS and coverslips mounted in Dako Fluorescent mounting medium (Agilent). For Drebrin1 staining, cells were fixed in cytoskeleton buffer supplemented with 4% PFA and 4% sucrose for 10 min at RT. The following antibodies and reagents were used: mouse monoclonal anti-HTT (4C8, IGBMC, Strasbourg), diluted 1:100; rat monoclonal anti-tyrosinated tubulin antibody (YL1/2, Provided by I Arnal, Grenoble Institute Neuroscience), diluted 1:4000; chicken anti-GFP (abcam), diluted 1:1000; Phalloidin-ATTO488 or phalloidin-ATTO647 (Sigma-Aldrich) to stain F-actin, diluted 1:1000; chicken anti-GFP (abcam) diluted 1:1000; anti-Drebrin1 (abcam) diluted 1:1000. Secondary antibodies were donkey anti-mouse IgG (H+L)-Cy™3 conjugate (Jackson ImmunoResearch), donkey anti-rat IgG (H+L)-Cy™5 conjugate (Jackson ImmunoResearch) and chicken IgY (H+L)-Alexa Fluor488 (Thermofisher).

### Confocal imaging

Growth cones were visualized with a confocal laser scanning microscope (Zeiss, LSM 710) and GaAsP detector (Zeiss Airyscan) using a ×63 oil-immersion objective (NA 1.4). The confocal stacks were deconvolved with AutoDeblur. To measure axonal length, we took large field images using an X20 objective (Zeiss, Axioscan or mosaic mode on Zeiss Confocal LSM710). Acquisitions were performed in the photo-imaging Facility of the Grenoble Institute of Neuroscience.

### Morphologic/Morphometric analyses

To assess axon length, we defined the axon as the neuron’s longest neurite, which had to measure at least 100 µm. (Lengths were measured using NeuronJ plugin in ImageJ software.) For branching analysis, we counted manually the number of axon primary branches that were at least 15 µm long and normalized them to the length of the axon bearing them. Only neurons having at least 1 branch were counted. For growth cone morphology analyses, filopodia were defined as actin filaments emerging at least 1 µm beyond the lamellipodial structure, which delimited the actin crown perimeter. Growth cone area was measured from the axonal shaft to the end of the area delimitated by phalloidin staining. Exploratory microtubules were measured from their tips to the main microtubule bundle in the central domain.

### Co-localization analysis of HTT with F-actin and microtubules in growth cones

Growth cones were fixed, immunostained and imaged as described above. To measure the distribution of HTT between F-actin and microtubules, we used an ImageJ macro generating masks on signals obtained for endogenous HTT, Tyrosinated tubulin, and Phalloidin on each stack. The number of 3D objects was measured using the 3D object counter plugin in Fiji software (https://imagej.nih.gov/ij/download.html). The number of objects detected for HTT staining overlapping with Tyrosinated tubulin or Phalloidin signals were quantified and expressed in percentage of the total number of HTT objects. Thresholds were adjusted manually.

### Growth cone fractionation

Growth cone and non-growth cone fractions were separated from P0 cortices as in (Leshchyns’ka and Sytnyk 2013). Briefly, twelve cortices were homogenized by 10 strokes using glass potters, loaded onto a discontinuous 0.75/1.0/2.33 M sucrose gradient and spin at 242,000 g for 1 h at 4 °C in a SW32Ti rotor (Beckman & Coutler). Growth cone fractions were collected at the interface load/0.75 M sucrose and non-growth cone fractions collected between 0.75/1.0 M sucrose and then analyzed by Western Blot.

### Permeabilization assay

At DIV4, cells were treated at 37 °C for 3 min with 10 µM Paclitaxel/Taxol (Sigma-Aldrich) in BRB80 buffer (80 mM PIPES, 1 mM MgCl_2,_ 1 mM EGTA, pH 6.7) that was supplemented or not with 0.05% Triton-X100 (Sigma-Aldrich) for the permeabilized or control conditions, respectively. After treatment, cells were immediately fixed as described above. Growth cones were immunostained for endogenous HTT and F-actin and imaged as described above, with the same parameters and the mean fluorescence intensities of Venus, HTT and Phalloidin signals measured within the growth cone.

### Microtubule preparation and co-sedimentation assay in lysate

Polymerization of porcine Tubulin (TEBU-BIO) was performed in BRB80 supplemented with GTP (1mM) and glycerol (2.5%) for 1h at 37 C. Paclitaxel/Taxol 50 mM (Sigma-Aldrich) was added for 15min at the end of polymerization. P0-P1 cortices were lysed at 50 mg/mL in BRB80 supplemented with protease and phosphatase inhibitors (Sigma-Aldrich), then subjected to vortex/ice cycles for 20 min. The lysates were cleaned by 245,000 g ultracentrifugation for 40 min at 4°C to discard potential polymerized microtubules and insoluble materials in the pellet. The supernatant was collected and incubated or not (control) with 2 µM of polymerized microtubules for 30 min at RT. The Tubulin detected by western blot in the control line is the endogenous Tubulin. Total fraction (input) was collected before ultracentrifugation at 100,000 g for 30 min at 23°C in a TLA100.3 rotors (Beckman & Coutler). Supernatant was discarded and pellet fractions were obtained by resuspending pellet in Laemmli buffer 1X with volume equal to the initial volume. 20 µl of each fraction were loaded onto the gel for western blot analysis.

### F-actin preparation and co-sedimentation assay in lysate

Purified rabbit skeletal G-actin (TEBU-BIO) was polymerized at 24 µM in actin polymerization (AP) buffer containing 20 mM imidazole, 100 mM KCl, 2 mM MgCl_2_, 0.5 mM ATP (Sigma aldrich), 1 mM EGTA, pH 7.0 for 45 min at RT. Then filaments were stabilized by addition of phalloidin-ATTO488 (Sigma-Aldrich) diluted 1:10 G-actin for 15 min at RT. P0-P1 cortices were lysed at 50 mg/mL in AP buffer supplemented with protease and phosphatase inhibitors (Sigma-Aldrich), then subjected to vortex/ice cycles for 20 min. The lysates were cleaned by 245,000 g ultracentrifugation for 40 min at 4°C to discard potential polymerized F-actin and insoluble materials in the pellet. The supernatant was collected and incubated or not (control) with 2 µM of polymerized actin filaments for 30 min at RT. The Tubulin detected by western blot in the control line is the endogenous actin. Total fraction (input) was collected before ultracentrifugation at 100,000 g for 30 min at 23°C in a TLA100.3 rotors (Beckman & Coutler). Supernatant was discarded and pellet fractions were obtained by resuspending pellet in Laemmli buffer 1X with volume equal to the initial volume. 20 µl of each fraction were loaded onto the gel for western blot analysis.

### Purification of FL-HTT and fragments

The expression and purification of full-length HTT have been described^28^. We cloned the HTT fragments, including B-C_HEAT, C_HEAT, and NTD1, with a modified pFastBac1 vector (Thermo Fisher Scientific) containing N-terminal FLAG tag, 6 x His tag, and a TEV cleavage site, and expressed them in Sf9 cells with Baculovirus. The purification method for B-C_HEAT, C_HEAT was identical to the method for full-length HTT, but the purification of NTD1 was distinct from the other two fragments. From harvesting the expressed cells to purifying them, the NTD1 must be contained within a high salt buffer condition (500 mM NaCl, 5% glycerol, and 50 mM Tris-HCl, pH8.0). NTD1 was therefore purified by affinity chromatography with Ni-NTA Agarose (QIAGEN), followed by cleavage of N-terminal tags with TEV protease. After recapturing the cleaved tags with Ni-NTA Agarose, the sample was purified with size-exclusion chromatography using HiLoad Superdex 200 26/600 (Cytiva) within 500 mM NaCl, 50 mM Tris-HCl, pH8.0. The HTT B-domain cloned into a pGEX-4T-1 vector was expressed with *Escherichia coli* BL21(DE3) (Thermo Fisher Scientific) at 18 °C in the presence of 0.5 mM IPTG for 18 h. The incubated cells were harvested with centrifugation (4,600 × g for 20 min) and re-suspended in buffer containing 100 mM NaCl, 50 mM Tris-HCl (pH 8.0), 5% glycerol. The suspended cells were lysed with sonication, and cell lysates were cleared by centrifugation at 27,000 × g for 1 hr. The GST-tagged Bridge domain was purified by affinity chromatography with glutathione-SepharoseTM 4B resins (Cytiva) followed by size exclusion chromatography with Superose 6 10/300 (GE Healthcare) in the running buffer containing 150 mM NaCl and 20 mM HEPES (pH 7.5).

### Co-sedimentation of F-actin with purified FL-HTT

A quantity of 3 µM of F-actin was incubated with 500 nM of FL-HTT in AP buffer for 30 min at RT and then endured high-speed centrifugation at 100,000 × g for 20 min at 23 °C. Supernatant and pellet were collected and loaded onto 8% SDS-PAGE for analysis after staining with Coomassie blue. The quantity of HTT found in the pellet was measured relative to the quantity of HTT found in the supernatant. To determine the apparent equilibrium dissociation constant Kd_app_, a constant quantity of 500 nM FL-HTT was incubated with increasing concentrations of F-actin for 30 min at RT and centrifuged at high speed. Then, supernatant and pellet were collected and loaded onto 8 % SDS-PAGE for analysis after staining with Coomassie blue. The quantity of FL-HTT found in the pellet and supernatant was expressed as % of FL-HTT found in the pellet. For determination of F-actin bundling, 2 µM F-actin were incubated with 150, 300, or 750 nM FL-HTT in AP buffer for 1 h at RT and the reaction was spun 15 min at low speed (15,000 × g) using a benchtop centrifuge to pellet bundles. The quantity of F-actin found in the pellet and F-actin found in the supernatant were expressed as % of F-actin found in the pellet. For observation of bundles by immunofluorescence, 2 µM F-actin were incubated in absence or presence of 750 nM FL-HTT for 1 h at RT. The reaction was stopped with BRB80 supplemented with 0.5 % glutaraldehyde and spun onto 12 mm coverslips in a centrifuge tube (Beckman & Coutler) containing BRB80 buffer using the SW32Ti rotor (Beckman & Coulter) at 100,000 × g for 30 min at 23°C. Coverslips were recovered and fixed cytoskeleton buffer supplemented with 0.5% glutaraldehyde, 0.1 % Triton-X100, and 10 % sucrose for 10 min at RT. Glutaraldehyde auto-fluorescence was quenched as described previously and coverslips were incubated with phalloidin-ATTO488, washed 3 times with PBS and mounted in Dako mounting medium before confocal imaging. For co-migration on glycerol gradient, F-actin and HTT fragments were incubated in a 6:1 molar ratio for 2 h at 4 °C. Samples were submitted to separation on a discontinuous glycerol gradient (30∼70 %) in BRB80 buffer (80 mM PIPES, 1 mM MgCl_2_, 1 mM EGTA, pH 6.7) at 100,000 × g for 2 h at 4°C. Fractions were collected, prepared and analyzed on SDS-PAGE.

### Transmission electron microscopy

A quantity of 2 µM of F-actin was incubated in the presence of 300 nM FL-HTT at RT for 15 min to quantify the proportion of tips bound by HTT, or for 1 h to visualize bundles. For negative staining, 4 μl of reaction solution were loaded onto freshly glow-discharged 400-mesh copper grid (Delta microscopies), blotted using filter paper (Whatman no. 4) and floated on 2% uranyl acetate in water. Grids were observed with a JEOL 1200EX transmission electron microscope at 80 kV. Images were acquired with a digital camera (Veleta, Olympus) at 150,000× magnification. Acquisitions were performed at the Electron Microscopy Facility of the Grenoble Institute of Neuroscience.

### TIRF microscopy

Chambers were prepared from glass slides and functionalized with silane-PEG (Creative PEGwork) as described previously^63^. Prior to actin polymerization experiments, chambers were incubated with a solution containing 0.1 mg/ml 1.5 kDa PLL-g-PEG (Jenkem) in 10 mM Hepes, pH 7.4 for 2 min at RT and washed with HKME buffer (10 mM Hepes, pH 7.4, 50 mM KCl, 5 mM MgCl_2_, 1 mM EGTA) containing 1% BSA. Polymerization of 1.5 µM actin mix containing 12% ATTO565 labelled G-actin and 88% unlabeled actin (Hypermol) was monitored in the presence of CT buffer, 125nM or 500 nM FL-HTT in HKME buffer containing 4 mM DTT, 0.1% BSA, 1 mg/ml glucose, 82 μg/ml catalase (Sigma-Aldrich), 0.58 mg/ml glucose oxidase (Sigma-Aldrich), 0.25% methylcellulose (Sigma-Aldrich) and 0.2 mM ATP. The time of the addition of CT buffer or FL-HTT to the actin mix was defined as t=0. Samples were observed on an inverted microscope (Eclipse Ti, Nikon) equipped with an Ilas2 TIRF system (Roper Scientific), a cooled charge-coupled device camera (EMCCD Evolve 512, Photometrics), a warm stage controller (LINKAM MC60), and controlled by MetaMorph software (version 7.7.5, Molecular Devices). Samples were excited with 491 nm laser light, and timelapse imaging (at 488 nm) was performed at 26 °C for actin polymerization, during 30 min at 1 frame per 10 s with a 100 ms exposure time. The analysis of the percentage of the actin network found in bundles was performed by using Icy and imageJ macro. In brief, TIRF images were enhanced for detection of filaments using the Feature detector (Jacob and Unser, 2004) module of Icy software (de Chaumont et al., 2012). For bundle determination, the maxima and standard deviation of intensity of 10 individual filaments were measured. Filament regions with an intensity over the mean maximal intensity of individual filament plus three standard deviations were considered as bundled region. After skeletization, the number of actin filament was determined. The percentage of actin bundling was calculated by dividing the length of bundled region by the length of total actin. For the measurement of the mean fluorescence intensity of bundles in order to approximate the number of filaments they contain, a linescan of 6 pixels wide was traced perpendicularly to the bundle axis and the maximum fluorescence intensity obtained for each bundle was plotted. Measurements were performed on the 10 brightest objects in the field at the indicated times.

### Migration in native conditions

E15.5 cortices were lysed in AP buffer, supplemented with proteases inhibitors and phosphatases inhibitors and resuspended in a Native sample buffer containing 250 mM Tris-HCl, 1% (p/v) Bromophenol blue, 25% glycerol. Lysate was loaded onto a 5 % native-PAGE. For purified protein, FL-HTT was incubated at the designated concentration in AP buffer and migration was performed in a 5 % native-PAGE. Gels were stained using Coomassie blue.

### MS-based proteomic characterization of F-actin associated proteins

Hdh^lox/lox^ cortical neurons were infected with a lentivirus expressing the Cre-recombinase LV-CRE (LV.pLV-EGFP-Cre, Addgene) at DIV0 to deplete endogenous HTT and then lysed at DIV4 in AP buffer supplemented with protease and phosphatase inhibitors. Lysate was submitted to a pre-clearing ultracentrifugation at 100,000 × g for 15 min at 4 °C. Then, lysate was added onto F-actin that was pre-incubated with CT buffer or FL-HTT for 1 h at RT to achieve final concentrations of 2 µM F-actin and 500 nM FL-HTT after addition of neuron lysate. After 15 min incubation of lysate with F-actin in absence or presence of FL-HTT at RT, samples were centrifuged at 100,000 × g for 15 min at 23°C, before discarding of the supernatant and resuspension of the pellet in Laemmli buffer. A control containing lysate only was added in order to determine which proteins are specifically pulled down with F-actin in the pellet. Four replicates were prepared for each condition. Samples were separated onto a homemade SDS-PAGE 10% and proteins stained with Coomassie blue. The very intense bands corresponding to exogenously added FL-HTT (∼350 kDa) and actin (∼42 kDa) were discarded from each lane before further preparation of the samples to improve MS detection of other proteins. Proteins were then in-gel digested using modified trypsin (sequencing purity, Promega), as previously described (Casabona et al., 2013). The resulting peptides were analyzed by online nanoliquid chromatography coupled to MS/MS (Ultimate 3000 RSLCnano and Q-Exactive HF, Thermo Fisher Scientific) using a 140 min gradient. For this purpose, the peptides were sampled on a precolumn (300 μm x 5 mm PepMap C18, Thermo Scientific) and separated in a 75 μm x 250 mm C18 column (Reprosil-Pur 120 C18-AQ, 1.9 μm, Dr. Maisch). The MS and MS/MS data were acquired by Xcalibur (Thermo Fisher Scientific).

Peptides and proteins were identified by Mascot (version 2.7.0.1, Matrix Science) through searches against the Uniprot database (*Mus musculus* taxonomy, September 2021 download), the sequence of human HTT, a homemade database containing the sequences of classic proteins found in proteomic analyses (human keratins, trypsin, etc.), and the corresponding reversed databases. Trypsin/P was chosen as the enzyme and two missed cleavages were allowed. Precursor and fragment mass error tolerances were set at 10 and 20 ppm, respectively. Peptide modifications allowed during the search were: Carbamidomethyl (C, fixed), Acetyl (Protein N-term, variable) and Oxidation (M, variable). The Proline software (Bouyssie et al. 2020) was used for the compilation, grouping, and filtering of the results (conservation of rank 1 peptides, peptide length ≥ 6 amino acids, peptide-spectrum-match score ≥ 25, allowing to reach a false discovery rate of peptide-spectrum-match identifications < 1% as calculated on peptide-spectrum-match scores by employing the reverse database strategy, and minimum of 1 specific peptide per identified protein group). Proline was then used to perform a compilation, grouping and MS1 label-free quantification of the identified protein groups based on specific and razor peptides. MS data have been deposited to the ProteomeXchange Consortium via the PRIDE partner repository (Perez-Riverol et al. 2019) with the dataset identifier PXD029990.

Statistical analysis of the MS-based proteomic dataset was performed using the ProStaR software (Wieczorek et al., 2017). We discarded proteins identified in the contaminant database, proteins identified by MS/MS in fewer than two replicates of one condition, and proteins detected in fewer than four replicates of one condition. After log_2_ transformation, extracted MS1 abundance values were normalized by Variance Stabilizing Normalization (vsn), before missing value imputation (slsa algorithm for partially observed values in the condition, and DetQuantile algorithm for totally absent values in the condition). Statistical testing was conducted with limma, whereby differentially expressed proteins were sorted out using a log_2_(Fold Change) cut-off of 1 and a p-value cut-off of 0.01, leading to an FDR inferior to 2% according to the Benjamini-Hochberg estimator.

The Gene Ontology term analysis was performed by the free software Gorilla, and enriched GO terms were chosen with a cut-off FDR at 1% (Eden et al., 2009). The list of proteins found stastistically enriched in the pellet with F-actin or FL-HTT+F-actin was compared to the list of the total proteins detected by MS. MS data have been deposited to the ProteomeXchange Consortium via the PRIDE partner repository with the dataset identifier PXD029990. MS data are available for reviewers at https://www.ebi.ac.uk/pride/login (Username; password: 58HIQdZ0).

### Statistical Analyses

GraphPad Prism (GraphPad Software, Inc.) software was used for statistical analyses. Outliers were identified by using ROUT method with Q=1% and removed for following statistical analysis. Normality of data distribution was assessed on control conditions by applying the d’Agostino-Pearson omnibus normality test. Parametric t-test was applied when data followed a normal distribution; the non-parametric t-test was used for non-normal data distribution. Paired or unpaired t-tests were applied.

## Acknowledgments

We thank C. Cuveillier and J. Delaroche for help with TIRF experiments and electron microscopy preparation, respectively; M. Sachs, and J. Beherens for providing the AMER2 plasmid; J. Courchet for providing Venus plasmid and helpful discussion; A. Antkowiak, C. Bosc, C. Fassier, A. Fourest-Lieuvin and V. Brandt for helpful discussions; the staff of the GIN animal, electron microscopy and Photonic Imaging Center facilities for advice and technical help. We acknowledge the contribution of AniRA lentivector production facility from the CELPHEDIA Infrastructure and SFR Biosciences (UAR3444/CNRS, US8/Inserm, ENS de Lyon, UCBL), especially G. Froment, D. Nègre and C. Costa. This work was supported by grants from Agence Nationale pour la Recherche (ANR-15-IDEX-02 NeuroCoG in the framework of the “Investissements d’avenir” program, SH; AXYON: ANR-18-CE16-0009-01, SH), Fondation pour la Recherche Médicale (FRM, équipe labellisée DEQ20170336752, SH; PhD fellowship, FDT202001010865, RC), AGEMED program from INSERM (SH) and by the National Research Foundation of Korea (NRF-2016K1A1A2912057 and NRF-2020R1A2B5B03001517 to JS) and by the Photonic Imaging Center of Grenoble Institute Neuroscience (Univ Grenoble Alpes – Inserm U1216) which is part of the ISdV core facility and certified by the IBiSA label. Proteomic experiments were partly supported by ANR under projects ProFI (Proteomics French Infrastructure, ANR-10-INBS-08) and GRAL, a program from the Chemistry Biology Health (CBH) Graduate School of University Grenoble Alpes (ANR-17-EURE-0003).

## Author contributions

RC, MC, IA and SH designed the study. RC and SH wrote the manuscript, which was commented on by all authors. RC performed most of the experiments. MC performed growth cone fractionation and contributed to cell biology experiments. HK performed protein purification and contributed to low- and high-speed centrifugation assays under the supervision of JS. JND performed the mass-spectrometry reading of samples under the supervision of YC. ED generated tools for image analyses. BB performed molecular biology experiments.

**Supplementary Information** accompanies this paper.

